# Linker histone regulates the myeloid versus lymphoid bifurcation of multipotent hematopoietic stem and progenitors

**DOI:** 10.1101/2024.09.16.613227

**Authors:** Kutay Karatepe, Bruna Mafra de Faria, Jian Zhang, Xinyue Chen, Hugo Pinto, Dmitry Fyodorov, Esen Sefik, Michael Willcockson, Richard Flavell, Arthur Skoultchi, Shangqin Guo

## Abstract

Myeloid-biased differentiation of multipotent hematopoietic stem and progenitor cells (HSPCs) occurs with aging or exhaustion. The molecular mechanism(s) responsible for this fate bias remain unclear. Here we report that linker histone regulates HSPC fate choice at the lymphoid versus myeloid bifurcation. HSPCs expressing H1.0 from a doxycycline (dox) inducible transgene favor the lymphoid fate, display strengthened nucleosome organization and reduced chromatin accessibility at genomic regions hosting key myeloid fate drivers. The transcription factor *Hlf* is located in one of such regions, where chromatin accessibility and gene expression is reduced in H1.0^high^ HSPCs. Furthermore, H1.0 protein in HSPCs decreases in an aspartyl protease dependent manner, a process enhanced in response to interferon alpha (IFNα) signaling. Aspartyl protease inhibitors preserve endogenous H1.0 levels and promote the lymphoid fate of wild type HSPCs. Thus, our work uncovers a point of intervention to mitigate myeloid skewed hematopoiesis.

## Introduction

Myeloid-skewed hematopoiesis underlies inflammation, cancer and other diseases^1,2^. Myeloid skewing can result from altered signaling between HSPCs and their bone marrow niche. For example, aged niches overproduce inflammatory cytokines such as interleukins and interferons (IFN), which by binding to their cognate receptors activate downstream effectors to promote myeloid differentiation^3–6^. Mutations in chromatin remodeling factors also could result in myeloid skewing, with myeloid-biased HSCs (my-HSCs) displaying pronounced epigenetic changes. For example, aged HSPCs display more open chromatin regions detectible by ATAC-seq^7^ and expanded H3K4me3 peaks detectible by ChIP-seq^8^. These results suggest that the myeloid versus lymphoid fate choice of HSPCs could have a chromatin basis, and aberrant chromatin opening may favor the myeloid fate. However, how chromatin changes lead to altered lineage potential of multipotent HSPCs remains unclear.

Eukaryotic nuclear DNA is packaged into a nucleoprotein complex as repeating nucleosome units, consisting of about 147 bp of DNA wrapped around an octamer of the core histones (H2A, H2B, H3 and H4). A fifth histone, the linker histone H1, associates dynamically with nucleosome core particles at the DNA dyad axis. H1 binding to chromatin stabilizes nucleosomes, increases chromatin folding/compaction, and is associated with a transcriptionally repressed state^9–11^. The function of linker histones has been perplexing because genetic inactivation of single H1 genes often failed to exhibit major phenotypes^12–14^, likely due to redundancy among the multiple H1 isoforms. Further reduction in H1 dosage by simultaneous inactivation of multiple H1 genes did result in profound phenotypes^15–17^, in association with derepressed expression of imprinted genes^18,19^ and repetitive elements^20^. Importantly, loss of silencing in interferon inducible genes has been seen as one of the consequences of low H1^21,22^. Overall, linker histones may regulate cell states by restricting chromatin opening at genomic regions important for certain cell types. Whether and how linker histone levels regulate HSPC chromatin organization and lineage fate decision has not been examined.

Within the hematopoietic system, low H1 level severely compromised lymphoid cells^16,17^, and bone marrow deficient in a single H1 gene (H1.0) appeared to form more myeloid colonies^23^. Therefore, we tested the hypothesis that the myeloid versus lymphoid fate choice of HSPCs could be sensitive to linker histone levels, modeled using a doxycycline inducible H1.0 transgene. We found that H1.0^high^ HSPCs favor the lymphoid fate, in association with strengthened nucleosome organization and repression of key myeloid driver genes. Furthermore, we show that the endogenous H1.0 level is amenable to physiologic and pharmacologic interventions. We propose a molecular model that could connect inflammatory signals to impaired lymphoid potential of HSPCs via the regulation of H1.0 levels.

## Results

### H1.0 overexpression in HSPCs leads to increased lymphopoiesis with expanded MPP4 and CLPs

As bone marrow deficient in H1.0 alone formed more myeloid colonies^23^, and aberrant H1.0 level has been implicated in several other pathophysiologic states^24,25^, we focused on modulating H1 dosage by expressing H1.0 as a transgene. We first expressed H1.0 as a GFP-fusion protein by lentivirus in wild type (WT) Lin-Sca1+ cKit+ (LSK) cells and transplanted them into irradiated syngeneic hosts. HMGN1-GFP and H2B-GFP were expressed as controls, with the former also binding to the nucleosome dyad and the latter widely used in HSPC tracing studies^26,27^. Following engraftment, GFP+ cells gave rise to distinct lineage contributions: H1.0-GFP+ cells yielded more lymphoid cells (both B220+ B and CD3+ T cells), whereas HMGN1-GFP+ cells had more myeloid progeny (CD11b+) (**Fig S1A,B**). The H2B-GFP+ cells gave rise to balanced lineages, consistent with the observation that H2B overexpressed as a fusion protein with the fluorescent Timer (FT) in HSPCs did not alter blood lineage distribution^28^. To better assess how H1.0 levels could regulate HSPC lineage choice, we generated knock-in mice by targeting the coding sequence for H1.0-GFP into the *Hprt* locus under a doxycycline (dox) inducible promoter^28,29^. Upon confirming dox-inducible transgene expression in the targeted mouse embryonic stem cells (mESCs) which express rtTA from the *Rosa26* locus, healthy and fertile mice containing the X^iH1.0-GFP^ allele (**Fig 1A)** were derived. Dox-dependent transgene expression in HSPCs was confirmed in LSK cells freshly sorted from iH1.0-GFP mice treated with dox in drinking water for 3 weeks (**Fig S1C).** An X^iHMGN1-mCherry^ allele was established in parallel as an additional control.

**Figure 1.**
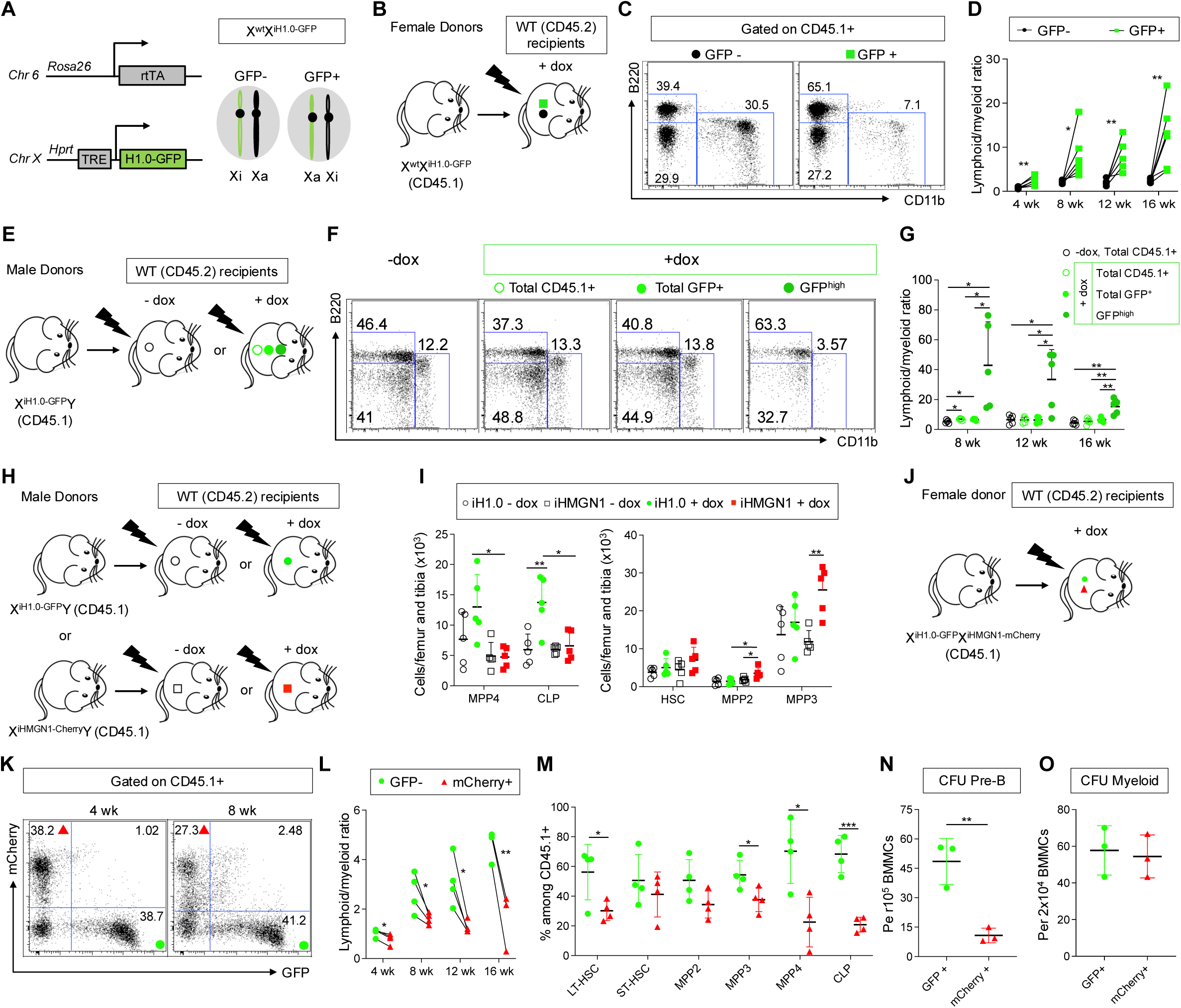
H1.0 expression in HSPCs confers increased lymphopoiesis. (A) Diagram for the transgene targeting strategy for introducing *H1.0-GFP* into the *Hprt* locus located on Chromosome X. *rtTA* is expressed from the *Rosa26* locus. Due to random X chromosome inactivation, heterozygous female cells could have the *iH1.0-GFP* transgene on either X chromosomes. The cells having *iH1.0-GFP* on the active X (Xa), but not those having it on the inactive X (Xi), will show dox-inducible transgene expression. (B) Scheme of the experimental setup for noncompetitive whole bone marrow (WBM) transplantation using X^wt^X^iH1.0-GFP^ heterozygous female mice as donors. All recipients are fed with dox water. (C) Representative FACS dot plots for the recipients in B. Donor-derived CD45.1+ cells in peripheral blood further divided into GFP- and GFP+ compartments. Representative lineage markers CD11b+ and B220+ cells at 16 weeks post-transplantation. (D) Quantification of lineage marker positive cells within donor-derived CD45.1+ cells divided into GFP- and GFP+ compartments as shown in C. Lymphoid/myeloid ratio is determined as (%B220+ + %CD3+)/%CD11b+ in individual recipient mice. n=6 each. (E) Scheme of the experimental setup for noncompetitive WBM transplantation using X^iH1.0-GFP^Y male mice as donors. The recipients are treated with regular or dox water. In the dox water treated recipients, donor (CD45.1+) cells are further divided by their H1.0-GFP fluorescence intensity. (F) Representative FACS dot plots for the recipients shown in E, at 16 weeks post-transplantation. – dox and + dox total CD45.1+, total GFP+ and GFP^high^ cells in peripheral blood stained for lineage markers (CD11b and B220) are shown. GFP^high^ denotes the top 10% among total GFP+ cells. (G) Quantification of the lymphoid/myeloid ratios in the recipients shown in F. (H) Scheme of the experimental setup for noncompetitive WBM transplantation using X^iH1.0-GFP^Y or X^iHMGN1-mCherry^Y male mice as donors. The recipients are treated with regular or dox water. (I) Quantification of the number of MPP4 and CLP per leg (femur and tibia) in recipient mice at 16 weeks post-transplantation shown in the left panel; the number of HSCs and MPP2 and MPP3 shown in the right panel. n = 5 per group. (J) Scheme of the experimental setup for noncompetitive WBM transplantation using X^iH1.0-GFP^X^iHMGN1-mCherry^ female mice as donors. All recipients are fed with dox water. (K) Representative FACS dot plots showing the donor derived (CD45.1+) peripheral blood cells at 4 and 8 weeks post-transplantation in recipients. (L) Quantification of the lymphoid/myeloid ratios within the respective GFP+ and mCherry+ cells within each individual recipient. n = 4 for 4-12 weeks; n = 3 for 16 weeks. (M) Quantification of bone marrow HSPC subsets positive for H1.0-GFP or HMGN1-mCherry in recipient mice at 12-16 weeks post-transplantation. n = 4 each. (N) Quantification of lymphoid colony-forming units by WBM cells at 16 weeks post-transplantation. n = 3 each. (O) Quantification of myeloid colony-forming units by WBM cells at 16 weeks post-transplantation. n = 3 each. Individual values as mean ± SD are shown. p<0.05, **p<0.01, ***p<0.001, by unpaired, 2-tailed Student’s t-test except for L, which was by paired, 2-tailed Student’s t-test. See also Figure S1.

As the *Hprt* locus is located on the X chromosome, dox-treated female heterozygotes (X^wt^X^transgene^) would contain a mixture of transgene+ and transgene-cells, due to random X inactivation. Expressed as fusion proteins, the fluorescence intensity could serve as a surrogate for transgene expression levels in live cells **(Fig 1A)**. We first confirmed the increased lymphoid cells from iH1.0-GFP+ HSPCs by transplanting X^wt^X^iH1.0-GFP^ whole bone marrow into lethally irradiated WT recipients and fed them dox water (**Fig 1B**). Donor (CD45.1+) cells fully reconstituted hematopoiesis in WT (CD45.2+) recipients (**Fig S1D**), and their contribution to B, T and myeloid lineages in the peripheral blood (PB) were monitored every 4 weeks for up to 16 weeks. Consistent with the lentivirally expressed H1.0, transgenic H1.0-GFP+ cells gave rise to increased lymphoid cells, a phenotype strengthening over time (**Fig 1C,D**). Despite the higher lymphoid reconstitution within the H1.0-GFP+ compartment, all recipients had complete blood cell counts within the normal range (**Fig S1E**). These results indicate that the H1.0-GFP+ cells follow homeostatic control together with the transgene-cells (**Fig 1A**). The increased lymphoid contribution by H1.0-GFP+ cells was also seen with male donors (**Fig 1E**), which was particularly prominent within the bright H1.0-GFP+ cells (**Fig 1E-G**). Furthermore and consistent with the lineage distribution observed by the virally expressed transgene (**Fig S1A,B**), mCherry+ cells derived from iHMGN1-mCherry donor bone marrow displayed decreased PB lymphoid:myeloid ratio, again in a transgene dosage dependent manner (**Fig S1F-H**). The opposing lineage effects following HMGN1 overexpression is consistent with the known H1-HMGN1 antagonism^30,31^ and the disturbed hematopoiesis in a Down syndrome model of HMGN1 overexpression^32^.

To assess the transgene effects on bone marrow HSPCs, we established cohorts of recipient mice engrafted with either X^iH1.0-GFP^Y or X^iHMGN1-mCherry^Y whole bone marrow, fed them with dox or regular water, and analyzed HSPC numbers 16 weeks later (**Fig 1H, S1I**). The lymphoid biased MPP4 and common lymphoid progenitor (CLP) were increased in mice engrafted with iH1.0-GFP marrow and fed with dox water (**Fig 1I**), while the myeloid biased MPP2/MPP3 and myeloid-committed progenitors increased in those engrafted with iHMGN1+ marrow (**Fig 1I, S1J**). The HSPC compartments in all control mice on regular water appeared similar and at an intermediate level (**Fig 1I**). Because the measurement of HSPC compartment sizes did not account for transgene expression levels, the modest difference between dox/regular water groups further supports a largely normal hematopoietic system from the transgene+ HSPCs. The larger differences between the two transgene+ groups is consistent with the interpretation that gain and loss of function in H1.0 drive opposing lineage effects, thereby increasing the dynamic range beyond that of WT control levels.

The X-chromosome hosted transgenes afford an opportunity to compare HSPCs expressing either transgene within the same animals, potentially reducing variability. Therefore, we noncompetitively transplanted whole bone marrow from female X^iH1.0-GFP^X^iHMGN1-mCherry^ donors into lethally irradiated WT recipients, and fed them dox water (**Fig 1J and S1K)**. While H1.0-GFP+ and HMGN1-mCherry+ cells were similar in abundance at 4 weeks (**Fig 1K and S1K,L**), %H1.0-GFP+ in all recipients increased over time **(Fig 1K)**, due to the increased B/T cells (**Fig S1M-N**), yielding higher lymphoid:myeloid ratios in each recipient (**Fig 1L**). 16 weeks after transplantation, the lymphoid-biased MPP4 as well as the lymphoid committed CLP expanded greatly in each recipient, with the iH1.0-GFP+ cells dominating these compartments (**Fig 1M**); in contrast, changes in MPP2/3 and myeloid-committed progenitors were minimal (**Fig 1M, S1O).** iH1.0-GFP+ bone marrow cells formed 5x more lymphoid colony forming units (CFU), with similar myeloid CFUs, than the iHMGN1-mCherry+ cells isolated from the same mice (**Fig 1N-O**). Taken together, elevating H1.0 level leads to increased lymphopoiesis starting at the MPP4-CLP stage and results in more B/T cells in the blood. Of note, the endogenous H1.0 mRNA is expressed at high levels in HSPCs and CLPs, such as documented by the *Tabula Muris* database^33^, due to H1.0 mRNA being polyadenylated (**Fig S1P**), supporting the notion that changes in H1.0 level accompany HSPC commitment to the lymphoid fate.

### H1.0-overexpressing HSPCs display reduced chromatin accessibility in subsets of genomic regions in association with strengthened nucleosome organization

Overexpressing H1 is expected to increase the H1/nucleosome ratio, better DNA protection by linker histones and correspondingly reduced chromatin accsibility^17,34^. To ascertain H1.0 protein overexpression in the transgenic HSPCs, we performed liquid chromatography mass spectrometry (LC-MS), using previously validated triple H1 knockout (TKO) mESCs as standard controls^17,19^. We found that H1.0 accounted for ∼2.2% of the total H1 repertoire in WT LSK cells, a value that increased to ∼11.5% in iH1.0+ LSK cells (**Fig 2A, Table S1**). The abundance of other major H1 isoforms was not significantly affected. Elevated H1.0 resulted in increased H1.0/nucleosome ratio, increasing it from 0.02 to 0.11 in LSK cells (**Fig 2B, Table S1**). Similar increase in H1.0 protein and H1.0/nucleosome ratio were detected in GMPs. These results confirm the elevated total H1.0 protein in the iH1.0-GFP+ HSPCs.

**Figure 2.**
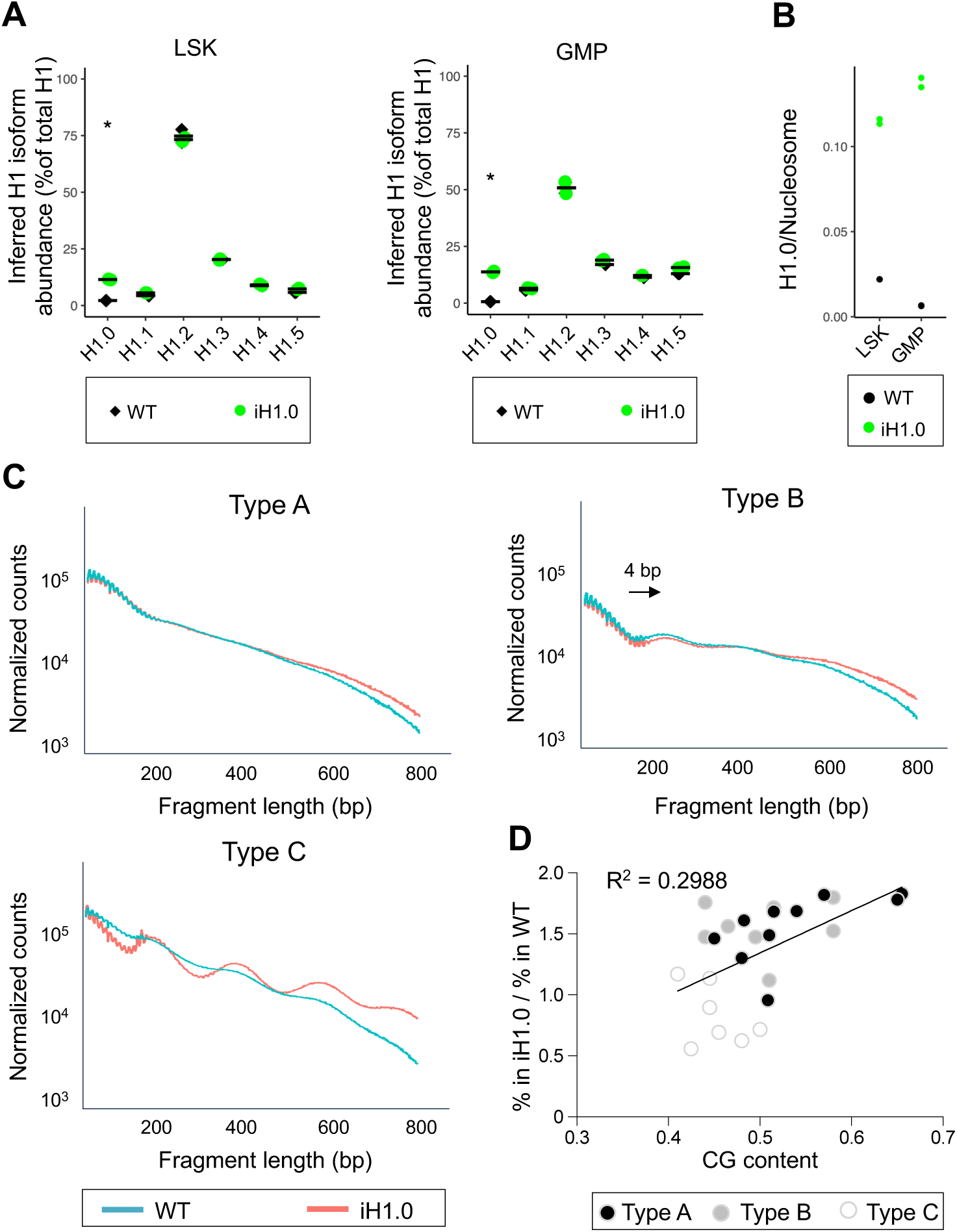
H1.0 overexpressing HSPCs display strengthened nucleosome organization. (A) Quantification of the inferred abundance of H1 subtypes as percent of total H1 by LC-MS in WT and iH1.0-GFP+ LSK and GMP cells. n = 2 each. (B) Quantification of the inferred H1.0/nucleosome ratio in WT and iH1.0-GFP+ LSK and GMP cells. (C) ATAC-seq fragment lengths and density in WT and iH1.0+ LSK cells. Three types of chromatin with regard to their nucleosome repeat signals are detected: Type A contains open chromatin and no nucleosome repeat signal in both WT and iH1.0 LSK cells; Type B has weak nucleosome repeat signal with an average of ∼4bp increase in nucleosome repeat length in iH1.0+ LSK cells; Type C has no nucleosome repeat signal in WT LSK but exhibits strong nucleosome repeats of 190 bp in iH1.0+ LSK cells. (D) Linear regression model depicting GC content vs. % of ATAC seq reads in each of the 25 annotated chromatin states in iH1.0+ relative to WT LSK cells. Reads coming from chromatin states with low GC content are underrepresented in iH1.0+ relative to WT LSK cells. Individual values as mean ± SD are shown. *p<0.05, **p<0.01, ***p<0.001, by unpaired, 2-tailed Student’s t-test. See also Figure S2.

We next assessed the nucleosome organization in the H1.0^high^ HSPCs. As nucleosomal DNA is less accessible to the Tn5 transposase, the enzyme used in ATAC-seq analysis, plotting the distribution of DNA fragment length in ATAC-seq data could inform about nucleosome organization^17^. We compared ATAC-seq results from iH1.0-GFP+ and WT LSK cells. Overall, the differentially accessible chromatin regions are more closed in iH1.0-GFP+ LSK cells, as expected from H1’s nucleosome compacting function (**Table S2**). With a cutoff of p < 0.001 and fold change of >4, we found 242 genomic regions being more closed and 78 regions more open in iH1.0+ LSK cells compared to WT LSK cells. Because nucleosome density is not uniform across the genome, with H1 enriched in heterochromatin and depleted on active chromatin regions^35^, we assessed the nucleosome repeat patterns across different chromatin states.

Following the 25 chromatin states defined using a combination of multiple chromatin markers^36^, we detected three types of chromatin according to their response to H1.0 overexpression (**Fig 2C and Fig S2**). Type A chromatin had minimal nucleosome repeat pattern in either WT or iH1.0+ LSK cells. Type B chromatin had weak nucleosome repeat signal in WT LSK and these nucleosome repeat signals increased in iH1.0+ LSK cells by ∼4 base pairs. The most significant change is seen in Type C chromatin, in which H1.0 overexpression substantially strengthened the nucleosome repeat pattern. Surprisingly however, Type C chromatin lacks obvious common features as it includes chromatin states ranging from quiescent (i.e. lacking all major chromatin markers tested) to heterochromatin to active chromatin^36^. Closer examination revealed that most Type C chromatin contains a lower percentage GC content (**Fig 2D**). Considering the previous report that H1.0 prefers to bind high GC regions^24^, our results suggest that the genomic regions disfavored by H1.0 in WT LSK gain nucleosome organization when H1.0 becomes abundant as H1.0 levels are increased by transgene expression. Taken together, these results suggest that the myeloid versus lymphoid fate choice in HSPCs could be sensitive to the strength of nucleosome organization in certain genomic regions where the properties of chromatin can be altered by elevating the level of a linker histone.

### H1.0 promotes HSPC lymphoid potential by reducing chromatin accessibility and gene expression of *Hlf*

Inspecting the genes located within the most differentially closed regions in iH1.0+ LSK cells revealed many known to be upregulated in myeloid biased HSCs^7,37^, such as *Slamf1* and *Itgb3* among others (**Table S2, Fig S3A**), suggesting that their elevated expression in myeloid biased HSCs could be related to chromatin de-repression due to reduced H1 level. In addition to these my-HSC marker genes, the genomic region encoding the transcription factor Hlf was among the top differentially closed regions in iH1.0+ LSK cells (**Table S2, Fig 3A**), most prominently at an intergenic region ∼1 kb 3’ distal to the *Hlf* gene. ATAC-qPCR with primers spanning this differentially closed region confirmed reduced chromatin accessibility in the iH1.0+ LSK cells (**Fig 3B**). This region and the *Hlf* gene body have ∼46% GC (**Fig S3B**). This differentially closed region is also gradually closed during WT HSC differentiation into the lymphoid biased MPP4 and almost completely lose ATAC-seq signals in CLP^7^ (**Fig 3A**). Together, these results suggest that chromatin accessibility at this region 3’-distal to *Hlf* is regulated during the lineage fate choices of WT HSPCs, despite the absence of annotated regulatory chromatin features.

**Figure 3.**
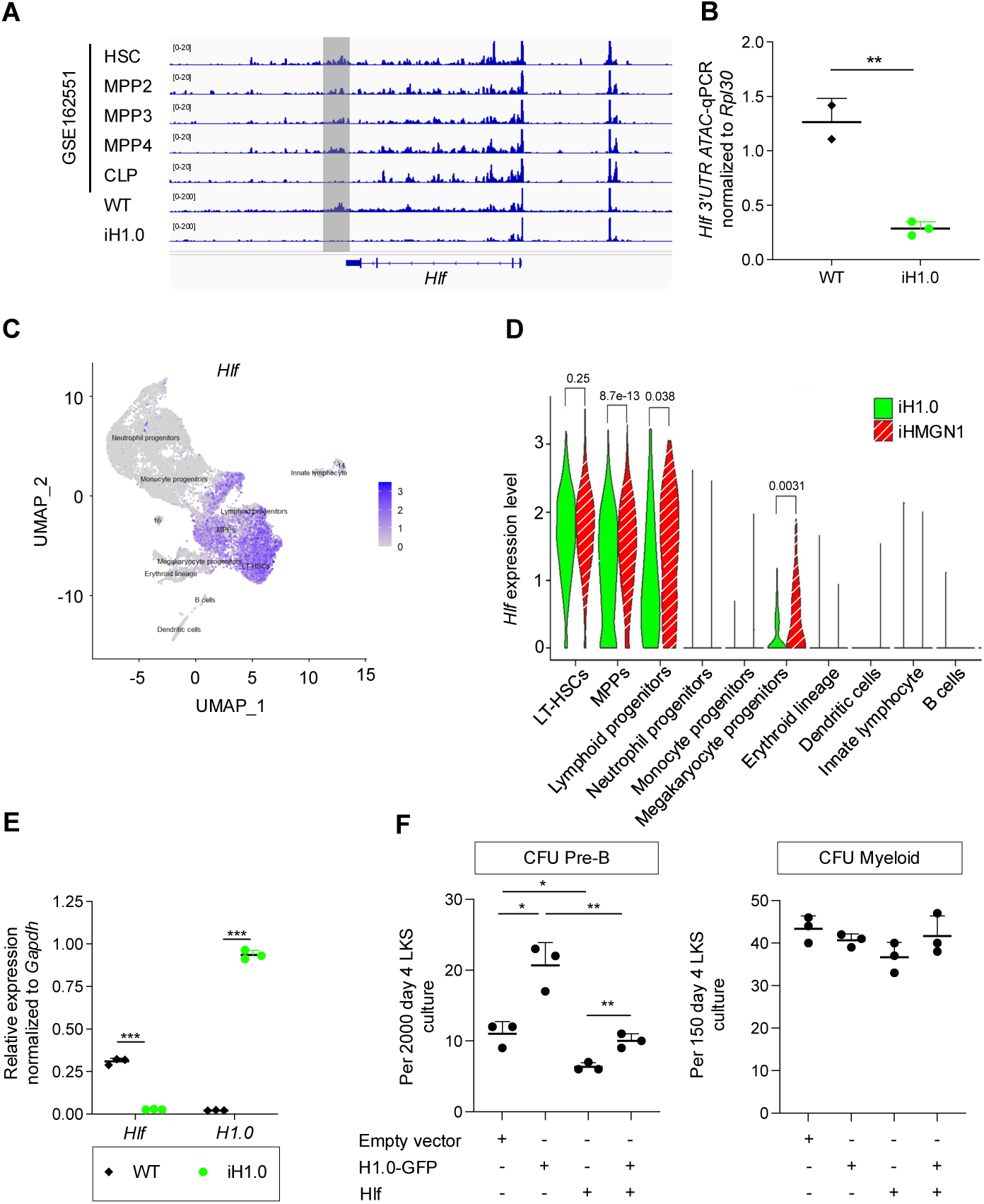
H1.0 expression imparts lymphoid fate potential by reducing chromatin accessibility and gene expression of *Hlf*. (A) Chromatin accessibility measured by ATAC-seq around the *Hlf* genomic region during WT HSPC differentiation from dataset GSE162551, aligned with those obtained from WT and iH1.0 LSK cells. n = 2 for WT and n = 3 for iH1.0. (B) ATAC qPCR quantification for the region highlighted in (A) from WT and iH1.0 LSK cells, normalized to that of mRPL30. n = 2 for WT and n = 3 for iH1.0. (C) HSPCs on UMAP plot of scRNA-seq of LSK cells from WT and X^iH1.0-GFP^X^iHMGN1-mCherry^ mice. Gene expression is plotted as log(normalized count + 1), with gray equal to no counts and dark blue representing the maximum value detected. Highest *Hlf* expression overlaps with the LT-HSCs. (D) Violin plots of *Hlf* mRNA levels across the cell types within iH1.0+ and iHMGN1+ LSK compartments. (E) *Hlf* and *H1.0* mRNA levels in freshly sorted WT and iH1.0+ LSK cells from mice on dox water for 3 weeks. N = 3 donors per group. (F) Quantification of lymphoid and myeloid CFU numbers using WT LSK cells transduced with indicated lentiviral constructs. n = 3 each. Individual values as mean ± SD are shown. *p<0.05, **p<0.01, ***p<0.001, by unpaired, 2-tailed Student’s t-test. See also Figure S3.

Hlf is a PAR bZip family transcription factor^38^ which is highly and specifically expressed in HSCs^39,40^. Hlf expression drives myeloid commitment while interfering with lymphoid differentiation^41^. Thus, the reduced chromatin accessibility at *Hlf* could potentially lead to its decreased expression and account for the superior lymphoid potential of H1.0^high^ HSPCs. Therefore, we assessed *Hlf* mRNA expression in HSPCs by single cell RNA-seq in LSK cells isolated from X^iH1.0-GFP^X^iHMGN1-mCherry^ mice treated with dox for 2 weeks. The co-expressed *GFP* or *mCherry* mRNA was used to distinguish the cells expressing either transgene (**Fig 3C-D**). WT LSK cells were analyzed as controls to identify lineage primed/committed progenitors according to established marker genes^42,43^ (**Fig 3C-D, S3C**). As expected, *Hlf* is highly expressed in LT-HSCs, and becomes downregulated as they differentiate into various progenitors. Intermediate to low levels of *Hlf* is detectable in lymphoid progenitors and megakaryocytic progenitors, but is undetectable in all the other cell types analyzed (**Fig 3C-D, S3C**). Corroborating H1.0 driven repression of *Hlf* expression, *Hlf* mRNA level was reduced in iH1.0+ MPPs and lymphoid progenitors, as compared to the iHMGN1+ counterparts, even though LT-HSCs of both genotypes had comparable *Hlf* (**Fig 3D**). Since all cells in these analyses were LSK, these results confirm *Hlf* expression to be highest in the multipotential HSPCs; within the cells whose *Hlf* expression is shutting down, abundant linker histones could facilitate its repressed expression. To confirm that H1.0 indeed represses *Hlf* expression, we compared the *Hlf* mRNA level by RT-qPCR in freshly isolated iH1.0+ and WT LSK and found that *Hlf* mRNA to be greatly reduced in H1.0+ cells (**Fig 3E**). Of note, *Hlf* expression became undetectable after 4 days of culture regardless of the genotypes (**Fig S3D**), consistent with the loss of *Hlf* expression as HSPC further differentiated. These data support that *Hlf* expression could be more effectively repressed in the presence of abundant H1.0 as the multipotent HSPCs commit lineage choices.

To determine whether the reduced *Hlf* expression in H1.0^high^ LSK cells is responsible for their higher lymphoid potential, WT LSK cells were transduced with viral constructs that overexpress either H1.0, Hlf or both, followed by plating into methylcellulose media permissive for either lymphoid or myeloid colonies (**Fig 3F**). H1.0 overexpression increased the lymphoid CFU, as expected. However, this increased lymphoid potential was abolished by Hlf co-expression. Similar myeloid CFU were formed in all conditions (**Fig 3F**). Taken together, these data support the model that high H1.0 promotes the lymphoid fate of multipotent HSPCs by reducing chromatin accessibility and gene expression of the myeloid driver gene *Hlf*.

### H1.0 level is amenable to physiologic and pharmacologic regulations

The results shown above suggest that H1.0 level might be a regulatory point to adjust the myeloid versus lymphoid lineage output. We reasoned that changes in H1.0 levels could be reflected by the fluorescence intensity of the H1.0-GFP fusion protein. In female X^iH1.0-GFP^X^iHMGN1-mCherry^ mice briefly treated with dox water (1 week) (**Fig 4A**), most HSPC compartments had roughly equal percentage of cells positive for either transgene, as expected from random X chromosome inactivation (**Fig 4B and Fig S4A**). However, the lymphoid-biased MPP4 was dominated by H1.0-GFP+ cells, yielding the apparent dominance by H1.0-GFP+ cells within the LSK cells. Strikingly, we noted that among LSK cells, the lymphoid-biased MPP4 also display brighter H1.0-GFP despite transgenic expression being driven by the dox-responsive promoter (**Fig 4B-C**). In contrast, HMGN1-mCherry intensity in MPP4 was similar to other LSK subsets (**Fig 4B and Fig S4B**). Longer treatment with dox water (3 weeks), during which H1.0-GFP+ HSPCs presumably underwent further differentiation, the CLPs displayed even brighter H1.0-GFP than the LSK **(Fig 4D-E)**. These results echo the expansion of MPP4-CLP following H1.0 overexpression (**Fig 1I, M**). Importantly, the lymphoid biased and lymphoid committed cells display brighter H1.0-GFP. As H1.0-GFP expression is driven by the transgenic promoter, we interpret the brighter H1.0-GFP fluorescence in MPP4-CLP to indicate potential regulation of H1.0 protein levels.

**Figure 4.**
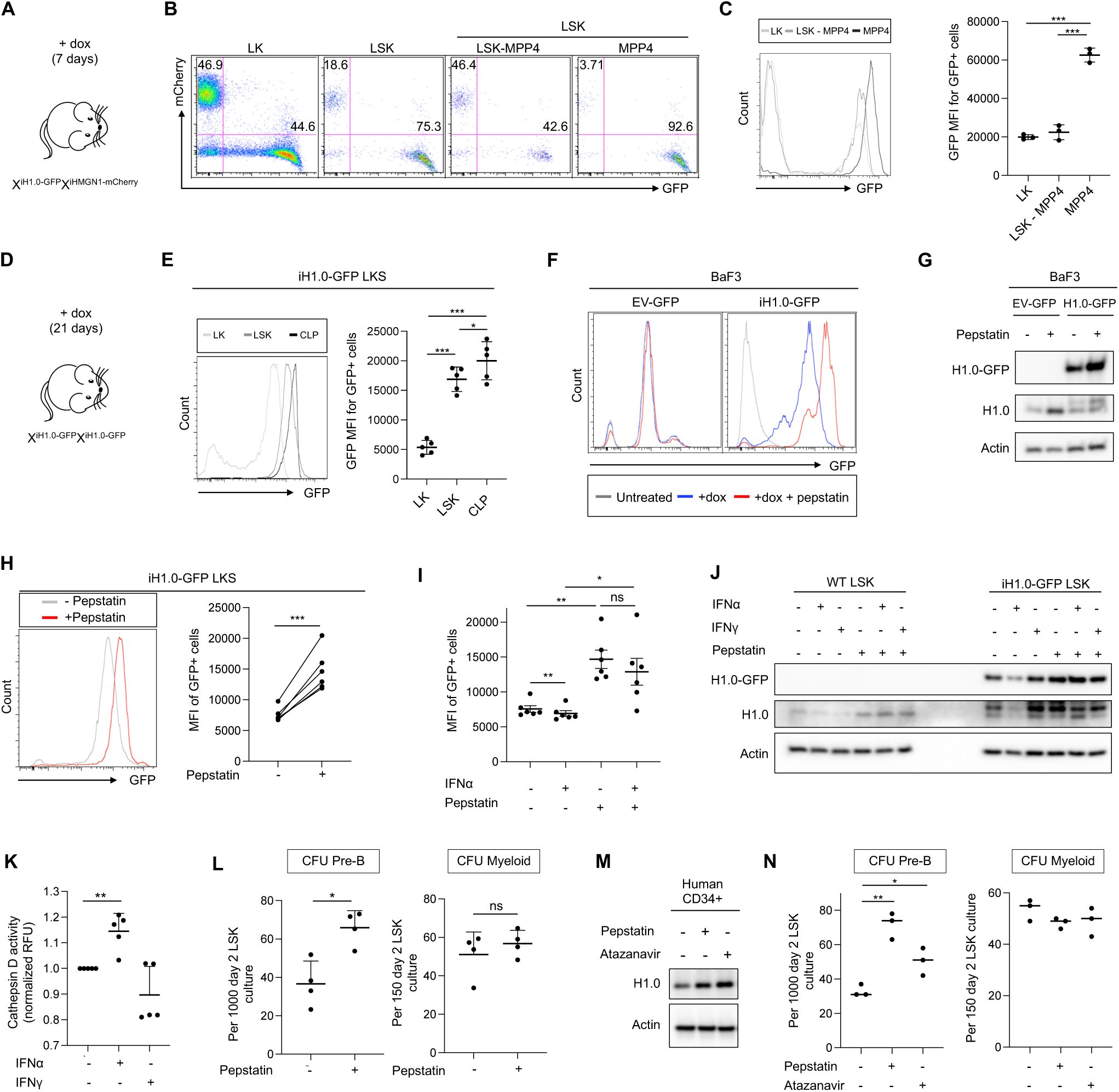
H1.0 level is amenable to physiologic and pharmacologic regulation. (A) Scheme of the experimental setup for analyzing bone marrow HSPCs from X^iH1.0-GFP^X^iHMGN1-mCherry^ female mice, which are unmanipulated except for one week of dox water treatment. (B) Representative FACS plots showing the distribution of LSK subsets that are positive for H1.0-GFP or HMGN1-mCherry in mice diagramed in A. n = 3 each. (C) Left: representative FACS histogram showing GFP fluorescence intensity in indicated LSK subsets shown in B. Right: quantification for GFP mean fluorescence intensity (MFI) for the GFP+ cells in indicated populations. n = 3 each. (D) Scheme of the experimental setup for analyzing bone marrow HSPCs from X^iH1.0-GFP^X^iH1.0^ ^GFP^ female mice, which are unmanipulated except for three weeks of dox water treatment. (E) Left: representative FACS histogram plot showing the GFP fluorescence intensity in indicated HSPC populations. Right: quantification for GFP mean fluorescence intensity (MFI) for the GFP+ cells in indicated populations. n = 5 each. (F) Representative FACS histogram showing GFP fluorescence intensity in BaF3 cells expressing lentiviral constructs: control pan-cellular GFP (EV-GFP) and dox inducible H1.0-GFP. Cells were cultured in the presence or absence of pepstatin. See also Fig S4C. (G) Representative Western blot using H1.0 and GFP antibodies in BaF3 cells expressing lentiviral EV-GFP or H1.0-GFP, in the presence or absence of pepstatin. Actin was probed as a loading control. (H) Representative FACS histogram plot showing GFP fluorescence intensity in freshly sorted iH1.0-GFP LSK cells cultured in the presence or absence of pepstatin. Right: quantification of GFP mean fluorescence intensity (MFI) in the GFP+ iH1.0 LSK cells. n = 6 each. (I) Quantification of GFP MFI in GFP+ iH1.0 LSK cells cultured in the presence or absence of IFNα and pepstatin. n = 6 each. (J) Western blot of endogenous H1.0 and H1.0-GFP in WT and iH1.0 LSK cells cultured in the presence or absence of IFNα/IFNψ +/-pepstatin. Actin was probed as a loading control. Results are representative of three independent experiments. (K) Cathepsin D activity as measured by a fluorescent substrate. Results are normalized to relative fluorescence units (RFU) in WT LSK cell lysates. n = 5 each. (L) Quantification of lymphoid and myeloid CFUs in LSK cells treated with pepstatin. Each dot is an independent experiment. n = 4 each. (M) Western blot of endogenous H1.0 protein in human CD34+ cells cultured in the presence of pepstatin or atazanavir. Actin was probes as a loading control. (N) Quantification of lymphoid and myeloid CFUs is LSK cells treated with indicated protease inhibitors. Each dot is a triplicate sample; results shown are representative of three independent experiments. n = 3. Individual values as mean ± SD are shown. *p<0.05, **p<0.01, ***p<0.001, by paired, 2-tailed Student’s t-test, except for 4N which is unpaired. See also Figure S4.

Given that proteases can regulate HSPC biology^44,45^, we hypothesized that some proteases may be involved in regulating H1.0 levels. We screened a panel of protease inhibitors for those that can increase H1.0-GFP intensity, using BaF3 cells expressing iH1.0-GFP from a retroviral construct. The H1.0-GFP+ BaF3 cells were treated with protease inhibitors, followed by flow cytometry assessment of H1.0-GFP fluorescence intensity (**Fig 4F, S4C)**. Inhibitors of cysteine proteases (E64), cysteine/serine/threonine protease inhibitor (Leupeptin) and calpains (calpeptin and PD150606) did not change the H1.0-GFP intensity; an inhibitor of serine proteases (pefabloc) had a mild effect, while the aspartyl protease inhibitor pepstatin consistently and significantly increased H1.0-GFP intensity (**Fig 4F, S4C).** Control BaF3 cells expressing the empty vector (EV) encoding GFP (EV-GFP) had similar fluorescence intensity across all treatment conditions. These results indicate that H1.0-GFP was reduced by aspartyl proteases. Importantly, pepstatin also increased the endogenous H1.0 protein in BaF3 cells (**Fig 4G**). We next examined whether pepstatin similarly regulates the H1.0-GFP level in LSK cells and found this to be the case (**Fig 4H)**. In contrast, pefabloc had negligible effects on H1.0-GFP intensity in LSK cells, although it had a mild effect in BaF3 cells (**Fig S4C-D**), and therefore was not pursued further. We reasoned that the changes in H1.0-GFP fluorescence intensity in LSK cells could serve as a reporter for identifying potential regulators of endogenous H1.0 as well as HSPC lineage fate choices.

To further explore the physiologic signals that regulate H1.0 levels, we wondered whether inflammatory signals could play a role, as inflammation often leads to myeloid biased hematopoiesis. We first treated H1.0-GFP+ LSK cells with a panel of inflammation inducers and measured their H1.0-GFP intensity (**Fig S4E**). We found that interferon alpha (IFNα) caused a reproducible reduction in H1.0-GFP intensity. Importantly, co-treatment with pepstatin blocked this reduction (**Fig 4I**). Next, we examined whether endogenous H1.0 levels are similarly regulated. As shown in **Fig 4J**, IFNα reduced the endogenous H1.0 level and this decrease was blocked by pepstatin even in WT LSK cells, paralleling the changes in transgenic H1.0-GFP in LSK cells. Consistent with IFNα induced aspartyl protease activation, we detected an increase in the proteolytic activity of one of the aspartyl proteases, cathepsin D, in LSK cell lysates (**Fig 4K**).

Lastly, we examined whether the changes in H1.0 level could be recapitulated by changes in HSPC lineage fate choices. Consistent with its H1.0-preserving effects, pepstatin increased lymphoid CFU numbers without changing the myeloid CFU of WT LSK cells (**Fig 4L**), indicating that pharmacologic inhibitors of aspartyl proteases could have lineage modulatory effects on HSPCs. Of note, the HIV protease is an aspartyl protease^46^, for which multiple safe and effective inhibitors have been in clinical use. To test the translational possibility that aspartyl protease inhibitors could modulate normal HSPC lineage fate choices, we treated human cord blood CD34+ cells with pepstatin or atazanavir, one of the clinically used HIV protease inhibitors. Both pepstatin and atazanavir increased the endogenous H1.0 protein level (**Fig 4M**). In agreement with the increased H1.0 level, similar to pepstatin, atazanavir also increased the lymphoid CFU without altering the myeloid CFU of WT LSK cells (**Fig 4N**). Taken together, our results implicate a potential translational opportunity for protease inhibitors to be repurposed to mitigate myeloid skewed hematopoiesis.

## Discussion

Our results depict a linker histone driven cell fate regulatory mechanism in HSPCs. We show that the multipotent HSPCs can be programmed to favor the lymphoid fate by increasing the level of a single linker histone, H1.0. These results contrast the observations that lack of single H1 genes is often of little phenotypic consequence^12,13,19^. It remains an open question whether elevating other H1 isoforms can lead to similar changes in HSPC lineage fate choices. Importantly, endogenous H1.0 level changes according to HSPC differentiation stage (**Fig S1P**) and/or by physiological signals, such as IFNα (**Fig 4J**). Linker histones have been reported to bind to interferon stimulated genes (ISGs) and suppress their expression^21^, whereas H1 depletion can trigger interferon response^22^. With our data showing that H1.0 level decreases in response to IFNα (**Fig 4J**), ISGs could be poised to hyperactivate when H1.0 is low, placing H1.0 and IFNα signaling in a positive feedback loop. This is consistent with the “inflamm-aging” concept where heightened/prolonged interferon signaling accompany aging and myeloid bias^47–50^. In such a model, aspartyl protease inhibitors could potentially break the vicious cycle consisting of inflammation and low H1, which appears to be attainable with pepstatin and a clinically used protease inhibitor atazanavir. Although histone proteolysis has been extensively studied in the formation of Neutrophil Extracellular Trap (NET) or NETosis^51,52^, proteolytic cleavage of core histones can occur without cell death as monocytes differentiate into macrophages^53^. Outside hematopoietic cells, cathepsin L has been reported to cleave H3 during mESC differentiation^54^ and cathepsin D cleaves H3 in the involuting mammary gland^55^. Defining the mode of H1 downregulation by proteases will substantially add to our understanding of how linker histones integrate with nucleosome and chromatin based cell fate control.

H1.0’s pro-lymphoid function is at least in part mediated by chromatin repression of the myeloid driver gene *Hlf*, whose cell type specific expression pattern may explain the cell context dependent response to H1.0 overexpression. Specifically, while LSK and GMP similarly express transgenic H1.0 (**Fig 2A-B**), the cell fate modulatory effect appeared to be much more pronounced in LSK subsets **(Fig 1M, S1J)** in association with H1.0 mediated repression of *Hlf* (**Fig 3**). It remains to be determined whether H1.0 can modulate fate choices in cells whose *Hlf* genomic region is already closed, such as in GMPs or via other target genes. Furthermore, the relationship between H1-based chromatin regulation and other epigenetic activities awaits further studies. For example, myeloid biased hematopoiesis due to Tet2 loss of function is common in both human and mouse HSPCs^56–58^; dissecting the role of H1 in the context of Tet2’s epigenetic dysfunction could provide further insights into the chromatin basis of HSPC fate regulation. The chromatin features that mediate much of the H1 response is surprising, as the most differentially closed chromatin region is 3’ distal to *Hlf* gene and lacks functional annotations (e.g. enhancer) – how nucleosome organization and chromatin accessibility at such regions crosstalk with regulatory mechanisms at the gene promoter/enhancers remain to be determined. Our data so far point to low GC% as a potential genomic feature in the differentially closed chromatin regions. In this regard, we note that Satb1, a protein that binds AT-rich genomic regions, promotes lymphopoiesis^59^, raising the possibility of functional cooperation between Satb1 and H1^60^, which also awaits future investigation.

## Acknowledgement

S.G and K.K were supported by DP2GM123507, R21DK128680 and U54 DK106857. K.K was partly supported by a U24 from U54 DK106829 and T32HL007974. B.M.F was supported by a Lo Graduate Fellowship and in part by the Coordenação de Aperfeiçoamento de Pessoal de Nível Superior - Brasil (CAPES). S.G. and B.M.F were supported by an ASH Bridge Grant. S.G was also supported by the Kutnick Family Foundation. H.P and A.I.S were supported by NIH grant R01GM147165 and D.V.F by NIH grant R01HD114814.

---

**Figure S1.**
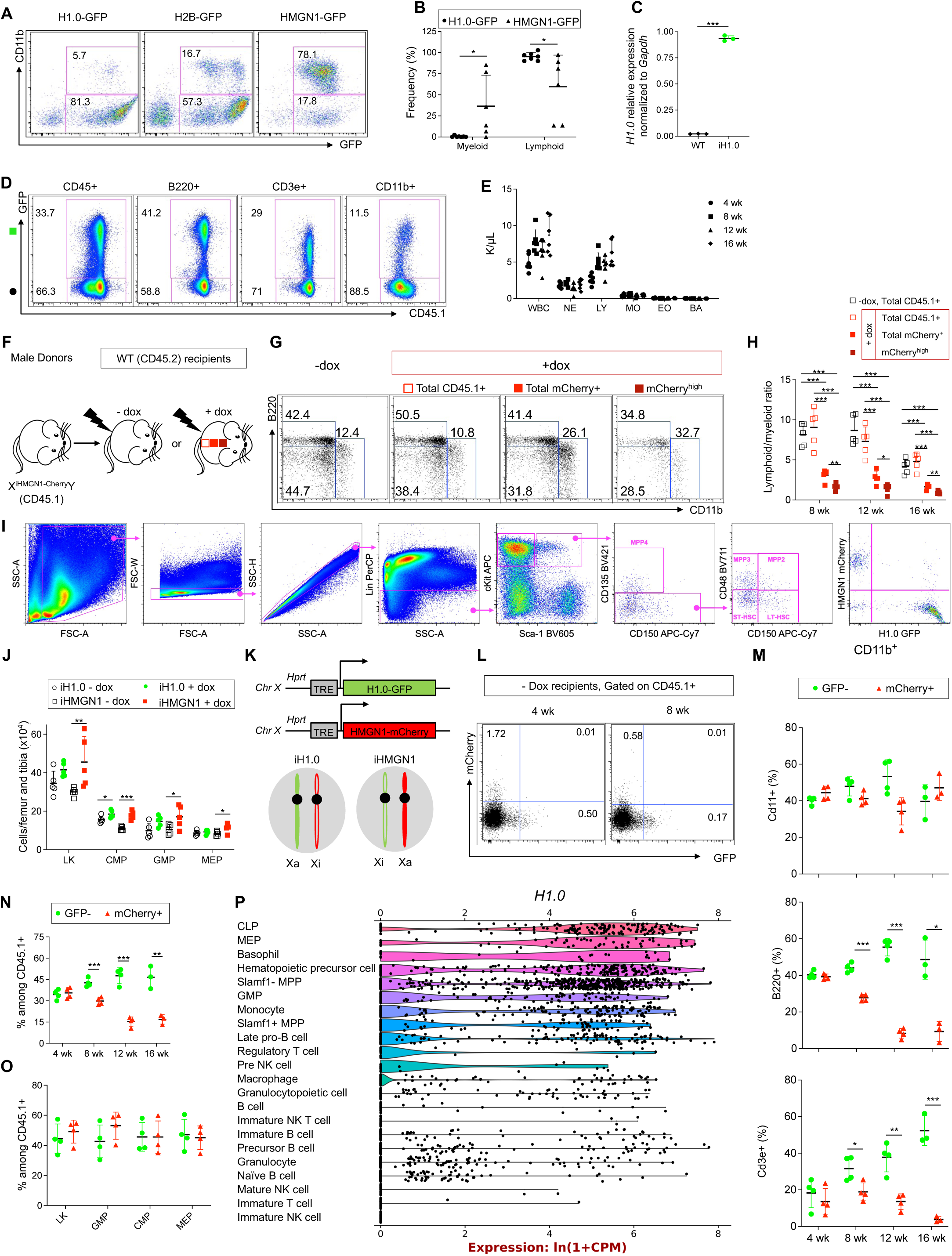
H1.0 expression in HSPCs confers increased lymphopoiesis *in vivo*. (A) Representative FACS dot plots for donor-derived peripheral blood cells at 8 weeks post-transplantation with LSK cells transduced with lentiviral H1.0-GFP, H2B-GFP and HMGN1-GFP. (B) Quantification of lineage marker+ cells within donor-derived GFP+ cells in recipients reconstituted with H1.0-GFP or HMGN1-GFP-transduced LSK cells at 16 weeks post-transplantation. n=6-7 per group. (C) *H1.0* mRNA level in freshly sorted LSK cells from WT (X^wt^X^wt^) and X^iH1.0-GFP^X^iH1.0-GFP^ mice treated with dox water for 3 weeks. n = 3 per group. (D) Related to Fig 1B-D. Representative FACS dot plots of peripheral blood B220+ B cells, CD3+ T cells and CD11b+ myeloid cells showing the GFP- and GFP+ cell percentage in the indicated lineage marker+ populations at 16 weeks post-transplantation. (E) Related to Fig 1B-D. Complete blood cell (CBC) counts including white blood cell, neutrophil, lymphocyte, monocyte, eosinophil and basophil cell counts of the WT recipients reconstituted with X^wt^X^iH1.0-GFP^ WBM 4-16 weeks post-transplantation. All recipients were maintained on dox water. n = 6 per group. (F) Scheme of the experimental setup for noncompetitive WBM transplantation using XiHMGN1-mCherryY male mice as donors. The recipients are treated with regular or dox water. In the dox water treated recipients, donor (CD45.1+) cells are further divided by their HMGN1-mCherry fluorescence intensity. (G) Representative FACS dot plots for the recipients shown in S1F, at 16 weeks post-transplantation. – dox and + dox total CD45.1+, total Cherry+ and Cherry^high^ cells in peripheral blood stained for lineage markers are shown. Cherry^high^ denotes the top 10% among total mCherry+ cells. (H) Quantification of the lymphoid/myeloid ratios in the recipients shown in S1G. (I) Representative FACS dot plots and gating strategy for HSPC subsets. (J) Related to Fig 1H-I. Quantification of the number of myeloid committed progenitors per leg (femur and tibia) in recipients at 16 weeks post-transplantation. n = 5 per group. (K) Related to Fig 1J. Diagram for comparing the H1.0-GFP+ and HMGN1-mCherry+ cells within the same recipients, engrafted with X^iH1.0-GFP^X^iHMGN1-mCherry^ female donor. When recipients are treated with dox water, all donor derived cells express either H1.0-GFP or HMGN1-mCherry, but not both, due to random X-chromosome inactivation. (L) Related to Fig 1K. Representative FACS dot plots showing the donor derived (CD45.1+) peripheral blood cells at 4 and 8 weeks post-transplantation in recipients that are on regular water. (M) Related to Fig 1L. Quantification for %H1.0-GFP+ and %mCherry+ cells among CD11b+ myeloid, B220+ B, CD3e+ T cells in the peripheral blood of recipient mice at 4, 8, 12 and 16 weeks post-transplantation. n = 4 for 4-12 weeks and n = 3 for 16 weeks. (N) Related to Fig 1L. Quantification for %H1.0-GFP+ and %mCherry+ cells among donor derived (CD45.1+) cells in peripheral blood of recipient mice at 4, 8, 12 and 16 weeks post-transplantation. n = 4 for 4-12 weeks and n = 3 for 16 weeks. (O) Related to Fig 1M. Quantification of bone marrow myeloid progenitor subsets positive for H1.0-GFP or HMGN1-mCherry in recipient mice at 12-16 weeks post-transplantation. n = 4 each. (P) Violin plot of single cell RNA-seq results depicting *H1.0* mRNA levels in bone marrow hematopoietic cells according to the Tabula Muris database. Individual values as well as means ± SD are shown. p<0.05, **p<0.01, ***p<0.001, by unpaired, 2-tailed Student’s t-test. See also Figure 1.

**Figure S2.**
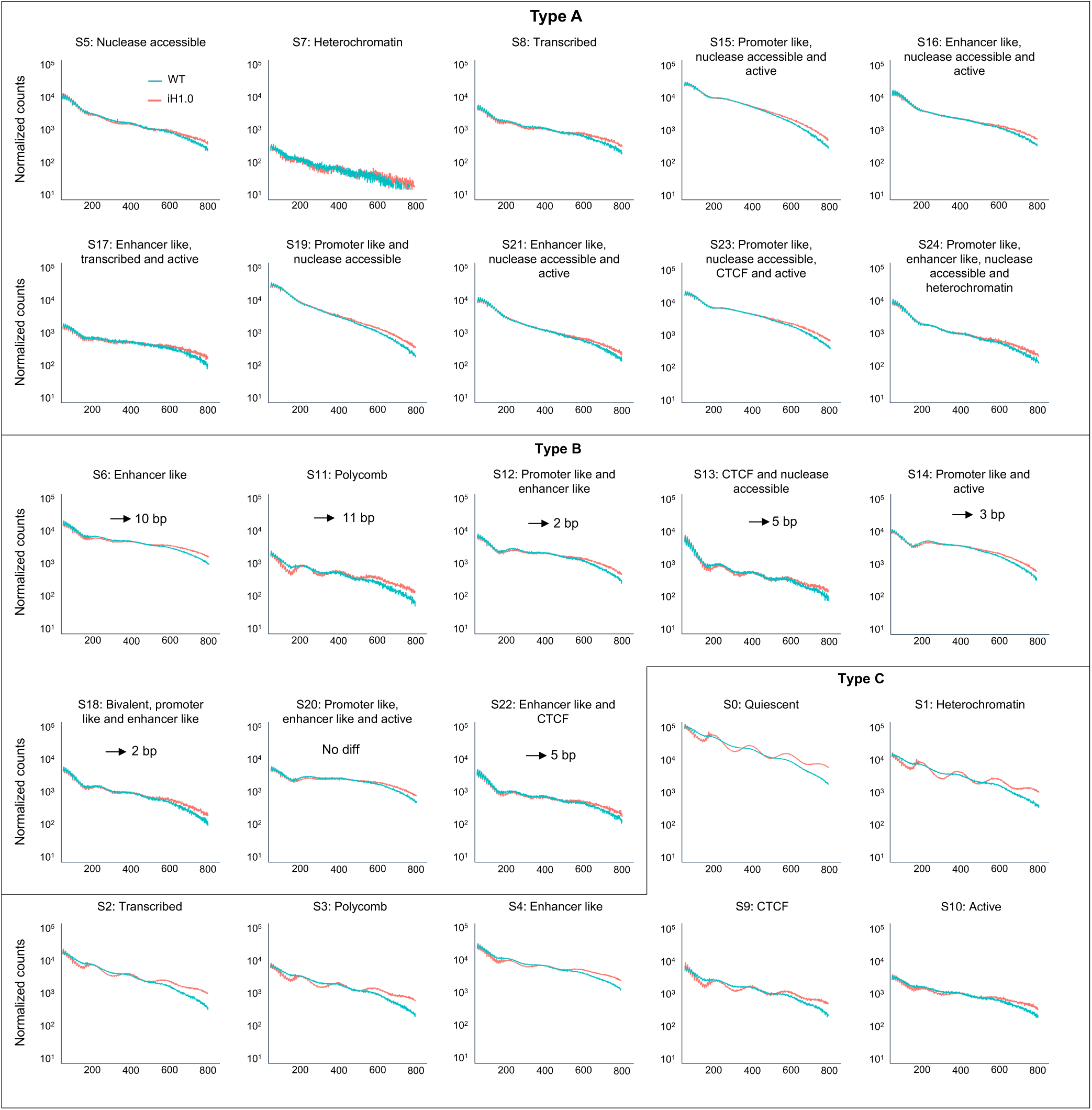
H1.0 overexpressing HSPCs display strengthened nucleosome organization. ATAC-seq fragment lengths and density in WT and iH1.0+ LSK cells across all 25 annotated chromatin state, as described by Xiang et. al. 2024. Change in nucleosome repeat length (right shift) was calculated using the “NRLfinder” as described by Willcockson et. al. 2021 and indicated on each plot with an arrow indicating the magnitude of change. Plots are grouped according to their behavior upon iH1.0 induction: unaffected (type A), right shift (type B), gaining nucleosome repeat signal with iH1.0 (type C). See also Figure 2.

**Figure S3.**
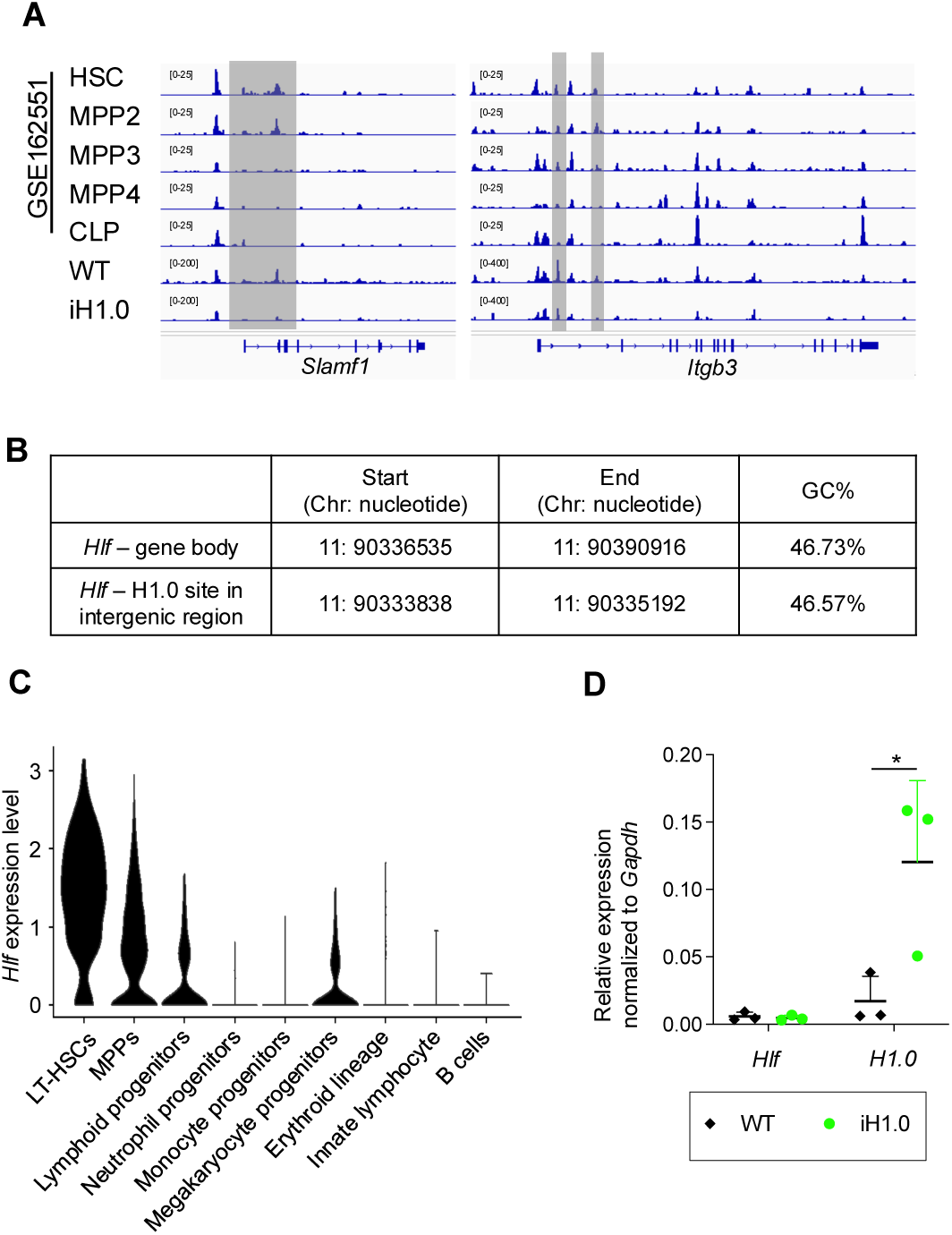
H1.0 expression imparts lymphoid fate potential by reducing chromatin accessibility and gene expression of *Hlf*. (A) Related to Fig 3A. Chromatin accessibility measured by ATAC-seq around the genomic regions of *Slamf1* and *Itgb3* during WT HSPC differentiation from dataset GSE162551, aligned with those obtained from WT and iH1.0 LSK cells. n = 2 for WT and n = 3 for iH1.0. (B) Related to Fig 3A, 2D. GC content of *Hlf* gene body and the differentially accessible intergenic region shown in Fig 3A. (C) Related to Fig 3D. Violin plots of *Hlf* levels across the cell types within WT LSK cells. (D) *Hlf* and *H1.0* mRNA levels in cultured WT and iH1.0 LSK cells. n = 3 donors per group. Individual values as well as means ± SD are shown. *p<0.05, **p<0.01, ***p<0.001, by unpaired, 2-tailed Student’s t-test. See also Figure 3.

**Figure S4.**
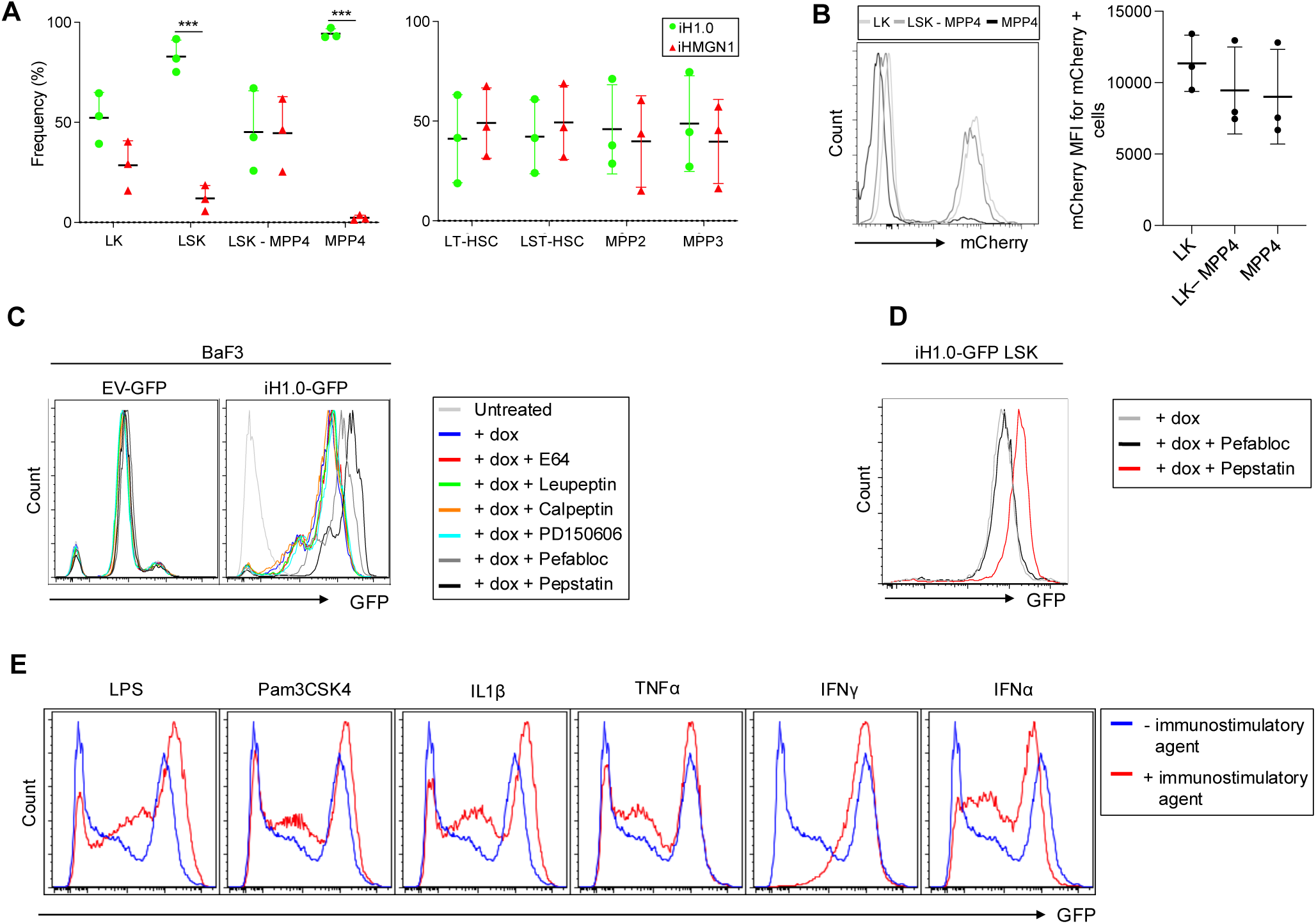
H1.0 level is amenable to physiologic and pharmacologic regulation. (A) Quantification for the distribution of LSK subsets that are positive for H1.0-GFP or HMGN1-mCherry from X^iH1.0-GFP^X^iHMGN1-mCherry^ female mice, which are unmanipulated except for one week of dox water treatment. n = 3 each. Related to Fig 4A-C. (B) Left: representative FACS histogram showing the mCherry fluorescence intensity in indicated HSPC populations. Right: quantification for mCherry mean fluorescence intensity (MFI) for mCherry+ cells in indicated populations. n = 3. (C) Representative FACS histogram showing GFP fluorescence intensity in BaF3 cells expressing lentiviral constructs: control pan-cellular GFP (EV-GFP) and dox inducible H1.0-GFP. Cells were cultured in the presence or absence of a panel of protease inhibitors including E64, leupeptin, calpeptin, PD150606, pefabloc and pepstatin. Pefabloc led to a mild increase in H1.0-GFP intensity. Results are representative of two independent experiments. See also Fig 4F. (D) Representative FACS histogram showing GFP fluorescence intensity in freshly sorted iH1.0-GFP LSK cells cultured in the presence or absence of pepstatin and pefabloc. Results are representative of six independent experiments. All cultures had dox added. (E) Representative FACS histogram showing GFP fluorescence intensity in freshly sorted iH1.0-GFP LSK cells cultured in the presence or absence of indicated immunostimulatory agent. All cultures had dox added. Results are representative of at least two independent experiments. Individual values as means ± SD are shown. *p<0.05, **p<0.01, ***p<0.001, by paired, 2-tailed Student’s t-test. See also Figure 4.

